# Extracellular vesicle-based vaccine platform displaying native viral envelope proteins elicits a robust anti-SARS-CoV-2 response in mice

**DOI:** 10.1101/2020.10.28.357137

**Authors:** K. Polak, N. Greze, M. Lachat, D. Merle, S. Chiumento, C. Bertrand-Gaday, B. Trentin, R. Z. Mamoun

## Abstract

Extracellular vesicles (EVs) emerge as essential mediators of intercellular communication. DNA vaccines encoding antigens presented on EVs efficiently induce T-cell responses and EV-based vaccines containing the Spike (S) proteins of Severe Acute Respiratory Syndrome Coronavirus (SARS-CoV) are highly immunogenic in mice. Thus, EVs may serve as vaccine platforms against emerging diseases, going beyond traditional strategies, with the antigen displayed identically to the original protein embedded in the viral membrane and presented as such to the immune system. Compared to their viral and pseudotyped counterparts, EV-based vaccines overcome many safety issues including pre-existing immunity against these vectors. Here, we applied our technology in natural EV’s engineering, to express the S proteins of SARS-CoV-2 embedded in the EVs, which mimic the virus with its fully native spikes. Immunizations with a two component CoVEVax vaccine, comprising DNA vector (DNA^S-EV^) primes, allowing *in situ* production of Spike harbouring EVs, and a boost using S-EVs produced in mammalian cells, trigger potent neutralizing and cellular responses in mice, in the absence of any adjuvants. CoVEVax would be the prototype of vaccines, where the sole exchange of the envelope proteins on EVs leads to the generation of new vaccine candidates against emerging viruses.

## Introduction

Extracellular vesicles (EVs) are membranous particles of endosomal origin and heterogenous size (30 to 150 nm) secreted by a vast majority of cell types from procaryotic to eucaryotic organisms. Embedded in their lipid bi-layer, transmembrane (TM) proteins and intra-luminal cargo serve as a go-between in cell-to-cell communication. EVs are capable of carrying antigenic information and triggering both humoral and cellular immune responses [1]. EVs carrying immunostimulatory molecules are already being explored for cancer immunotherapy and evaluated for other applications in multiple clinical trials [2]. Importantly, mammalian cells infected with viral pathogens, including human cytomegalovirus (CMV) and Human Immunodeficiency Virus (HIV), can release “antigenic” EVs containing viral components and triggering immune responses [3, 4]. EV-bound antigens released after DNA vaccination are more immunogenic and trigger stronger cytotoxic T-cell responses when compared to soluble antigens in murine tumour models [5]. It has been demonstrated that the Spike (S) envelope protein, presented on EVs, can trigger a humoral response against Severe Acute Respiratory Syndrome Coronavirus (SARS-CoV) which caused the SARS outbreak between 2002 and 2004 [6].

To exploit EVs as natural vaccines, we have developed a technology that allows loading of EVs with any fully native membrane protein [7]. In brief, a protein of interest is fused to a proprietary Pilot Peptide (CilPP), which interacts with the Endosomal Sorting Complexes Required for Transport (ESCRT) cellular machinery to sort the resulting chimeric protein onto EVs. As a response to the ongoing Coronavirus Disease 2019 (COVID-19) pandemic, efforts multiplied to develop vaccines against SARS-CoV-2 virus. We took advantage of this EV based platform to develop a non-adjuvanted, virus free vaccine against SARS-CoV-2, with native Spike protein loaded onto extracellular vesicles.

The Spike envelope protein of coronaviruses is synthesized in an immature form cleaved into S1 and S2 mature proteins. This maturation of S1 and S2 proteins is a prerequisite for the Spike functionality during cell infection. Majority of the tested vaccine candidates are based on the extracellular or the Receptor Binding Domain (RBD) of the Spike leaving behind potential T- and B-cell epitopes of the well conserved S2 protein [8]. Other enveloped viruses, including HIV and influenza also possess type I transmembrane envelope proteins which enable their cellular entry. Similarly to coronaviruses, these proteins are synthesized as trimers that are then cleaved into two mature proteins equivalent to S1/S2 (SU/TM for HIV and HA1/HA2 for influenza viruses), forming a “head” and a “stem” region, the latter anchoring the mature trimer into the viral membrane and being responsible for the virus-cell membrane fusion. The immunogenicity of viral envelope proteins is decreased by surface glycosylation and by the trimeric structure of the spike, which occludes important epitopes. Decades on HIV vaccine development revealed an absolute necessity for using a fully native envelope protein embedded in a membrane in order to obtain a high quality immune response. Indeed, HIV envelope SU head subunit vaccine provided no protection against infection in human, while triggering neutralizing antibody production in small animals and non-human primates [9]. Additionally, influenza neutralizing antibodies targeting HA1 head epitopes are usually susceptible to strain variability [10]. Importantly, all stem domains (S2, TM, or HA2) are more conserved than head domains (S1, SU and HA1). In HIV, the stem contains a highly conserved Membrane Proximal External Region (MPER) potent neutralizing epitope [11]. At the same time, HA2 stem is the target of broadly neutralizing antibodies [12]. Thus, vaccines that mount broad protection against either influenza, HIV or other enveloped viruses must elicit anti-stem antibodies [13]. Consequently, chimeric viruses or virus-like particles (VLPs) harbouring native envelope proteins seem the best platforms considering the conformation of a potential MPER peptide and the scaffold feature of lipids [14].

In addition to a humoral response, a consensus arises that triggering a strong T-cell immunity, even in the absence of cross-reactive neutralizing antibodies correlates with a protection against several viral strains [15, 16]. Work on SARS-CoV and Middle East Respiratory Syndrome Coronavirus (MERS-CoV) already revealed the importance of both humoral and cellular immunity in inducing a protective response [17, 18]. Finally, there is mounting evidence for a positive correlation between the resistance to the virus, the level of T-cell responses in SARS-CoV-2 infected people and T-cell cross-reactivity against S2 [19, 20].

## Materials and Methods

### Expression vectors and molecular cloning

Expression system to sort membrane proteins on extracellular vesicles (EVs) was developed by Ciloa SAS (Patent WO2009115561) [21]. Briefly, gene encoding the Spike from SARS-CoV-2 Wuhan-Hu-1 isolate (MN908947.3) was codon-optimized for human cells and cloned into inhouse eukaryotic expression vector [7] in order to generate the DNA^S-EV^ vector. SARS-CoV-2 Spike was fused in C-terminus to a pilot peptide (designated here as CilPP) which sorts the Spike protein to the surface of EVs. Modifications were introduced including the replacement of the spike glycoprotein signal peptide by that of metabotropic glutamate receptor 5, two consecutive proline substitutions at amino acid positions 986 and 987 and K776A/Q779G substitutions. Using the DNA^S-EV^ construct, the S-Trim expression vector was generated (DNA^S-Trim^) by the deletion of S2 domain downstream of the glutamine residue 1208 and by merging this truncated S2 with a bacteriophage T4 foldon trimerization motif followed by 6 Histidines [22]. To establish pseudovirus neutralization assay, the following constructs were obtained. The SARS-CoV-2 Spike gene from plasmid pUC57-2019-nCoV-S-Hu (GenScript, Piscataway, USA) was truncated in its cytoplasmic tail of 19 amino acids, in order to enhance pseudotyping efficiency [23], and then subcloned into in-house expression vector. We used the pNL4-3.Luc.R-E-luciferase reporter vector from NIH-AIDS Reagent Program [24] and exchanged the luciferase gene by the Nano Luciferase gene (named hereafter pNL4-3.NanoLuc). Both Homo sapiens angiotensin I converting enzyme 2 (ACE2) gene (NM_021804.2) and Homo sapiens transmembrane serine protease 2 (TMPRSS2) gene (NM_005656.4) cloned into pcDNA3.1(+), were from GenScript (GenScript, Piscataway, USA).

### Production and purification of SARS-CoV-2 protein S trimer (S-Trim)

The purification of coronavirus S Trimer was described earlier [22]. In brief, HEK293T cells were transfected with DNA^S-Trim^ plasmid using PEI, harvested 48h after, and S-Trim protein was purified in a buffer containing 0.5 mM PMSF, Protease Inhibitors (Protease Inhibitor Cocktail set III, EDTA-free, Calbiochem), 2% Octyl β-D-1 thioglucopyranoside, 12 mM MgCl_2_ and DNase I. After a centrifugation at 800 rpm for 10 minutes at 4 °C and then at 15 000 rpm for 15 minutes at 4°C, the supernatant was collected, equilibrated with 10 mM Imidazole, and purified over IMAC resin (HisTALON™ Superflow Cartridge, TAKARA), using a fast protein liquid chromatography system (ÄKTA FPLC Systems, GE Healthcare). S-Trim was eluted with 150 mM Imidazole. Fractions containing pure S-Trim were identified by Western Blot and pooled. The protein concentration was estimated with BCA assay (Pierce BCA Protein Assay Kit, ThermoFisher Scientific) by reading the absorbance at 562 nm.

### Production of EVs harbouring SARS-CoV-2 spikes in mammalian cells

HEK293T were cultured in DMEM supplemented with 5% heat inactivated foetal bovine serum (iFBS), 2 mM GlutaMAX and 5 μg/mL of gentamicin at 37 °C in a 5% CO_2_ humidified incubator. HEK293T cells were routinely tested and found negative by MycoAlert™ mycoplasma detection kit (Lonza Nottingham, Ltd.). HEK293T cells were plated into cell chambers of 10 trays in one litre of complete medium and were transfected with DNA^S-EV^ expression plasmid using PEI. Twenty-four hours post-transfection, cultures were fed with a medium supplemented with EV-free iFBS and incubated for a further 48 hours.

### EV purification

Cell culture medium was harvested from transiently transfected HEK293T cells and the EV isolation was performed as previously described [7, 25]. Briefly, cell culture supernatant was clarified by two consecutive centrifugations: 10 minutes at 1300 rpm and 15 minutes at 4000 rpm, both at 4 °C, followed by filtration through 0.22 μm membrane filters. The supernatant was then concentrated by ultra-filtration/diafiltration and purified by Size Exclusion Chromatography (SEC). Fractions containing EV markers (CD81 and CD63) were identified by ELISA. EV fractions containing Spike protein identified by WB were pooled, concentrated when necessary and used for analysis and injections.

### ExoView EV characterization

Purified EVs were sized and characterized using the Single Particle Interferometric Reflectance Imaging Sensor (SP-IRIS) technology using the ExoView R100 platform (NanoView Biosciences, Boston, USA). Tetraspanin kit chips (EV-TETRA-C), with capture antibodies against CD81, CD63, CD9 and control IgG (MIgG), were used to detect EVs. EVs (10^10^ EVs/ml) were coated on chips, then probed with detection antibodies provided with the kit (CD81/CF55, CD63/CF647, CD9 CF488A) according to the manufacturer’s protocol. The chips were imaged with ExoView R100, label-free counts were used for EV size determination (with a size range set from 50 to 200 nm).

### SDS-PAGE, Western blotting and antibodies

Protein concentration of EVs was measured using the BCA assay (Pierce BCA Protein Assay Kit, ThermoFisher Scientific). EVs or S-Trim preparations were separated by SDS-PAGE on a 4−15% acrylamide gel (4−15% Mini-PROTEAN® TGX Stain-Free™ Gel, Bio-Rad) and subsequently transferred onto PVDF membrane. For WBs in non-reducing conditions, a loading buffer without DTT was used. The immunodetection of SARS-CoV-2 spike protein was performed with primary antibodies against either S1 domain (SARS-CoV-2 Spike S1, rabbit monoclonal antibody [HL6], GTX635654, Genetex), or S2 domain (SARS-CoV-2 Spike S2, mouse monoclonal antibody [1A9], GTX632604, Genetex), or CilPP (in-house antibody raised in rabbit). Membranes were then incubated with the corresponding secondary HRP-conjugated antibodies (donkey anti-mouse or anti-rabbit HRP, #715-035-150 or #711-035-152 from Jackson ImmunoResearch); the signals were detected using an enhanced chemiluminescence detection kit (Super Signal West Pico Plus, 34580, ThermoFischer Scientific) and membranes imaged with ChemiDoc Imaging System (Bio-Rad). The same anti-S1 and anti-S2 antibodies as well secondary antibodies were also used to detect the Spike protein on EVs by ELISA (see protocol below).

### Ethics statement

All animal studies were performed in accordance with national regulations and approved prior to experimentation by the Ethics Committee for Animal Testing Languedoc-Roussillon (national agreement CEEA-036) and French Ministry of Research with the reference APAFIS #24869 (10 April 2020). All the procedures and protocols were performed at DMEM unit of INRAE-UM in Montpellier, France (Veterinary Services National Agreement n° E34-172-10, 04 March 2019).

### Mice and study design

Female BALB/cAnNCrl mice, 6 weeks old, were purchased from Charles River-Italy and placed into four groups of n=6. Mice were housed in individually ventilated cages with controlled environmental parameters: 24 degrees, 12 hours/12 hours light and dark cycle, nesting cotton squares for enrichment, free access to standard A04 irradiated food (SAFE, R04-25) and tap water. Behaviour and health status were observed daily and weight checked weekly. One week after their arrival, animals received 2 prime immunizations at day 0 and 21 and a boost immunization at day 42. Accordingly, Group 1 (named DNA^S-EV^/S-EV) received two DNA^S-EV^ injections and one S-EV injection. Group 2 (named DNA^S-EV^/S-Trim) received two DNA^S-EV^ injections and one S-Trim injection. Group 3 received three S-EV injections. Group 4 received three injections with PBS. DNA^S-EV^ vectors were injected using a Gene Gun (Bio-Rad, Helios) into the abdomen of mice previously shaved. Each mouse received 3 cartridges coated with gold beads containing DNA ^S-EV^ corresponding to a total of 3 μg of DNA. S-EVs were injected subcutaneously, 10 μg per mouse in 100 μl of PBS. S-Trim was injected subcutaneously in the same volume. Sera were collected via submandibular bleeding before each immunization: 70 to 100 μl of blood was collected with GOLDENROD animal Lancets (3mm, Genobios). All animals were euthanized at day 63 and spleens were collected for cellular response analysis.

### S1/S2 and antigen specific IgG ELISA

SARS-CoV-2 specific antibody titers of sera were determined by ELISA. Briefly, MaxiSorp ELISA plates (Nunc) were coated with 2 μg/ml of SARS-CoV-2 Spike overlapping peptide pools corresponding to whole S1 or S2 proteins (JPT, #PM-WCPV-S) in 100 μl 50 mM sodium carbonate/bicarbonate pH 9.6 buffer per well, overnight at 4 °C. Coated plates were washed 3 times with 200 μl of 1X PBS and saturated 1h at 37 °C with 200 μl 3% BSA in 1X PBS per well. Plates were washed three times with 1X PBS, and incubated in 3% BSA and 5% FBS with 3-fold mouse sera dilutions (starting from 1:100) for 2h at 37 °C. This was followed by 3 washes with 200 μl of 1X PBS per well and incubation with 100 μl per well of secondary donkey anti-mouse HRP conjugated antibody diluted 1:10,000 in 3% BSA in 1X PBS. Following the incubation with the secondary antibody, plates were washed 5 times with 200 μl of 1X PBS per well and developed with 100 μl of TMB per well (Bio-Rad, #R8/R9) for 30 minutes. The reaction was stopped by adding 50 μl of stop solution (2N sulfuric acid) per well. The 450 nm absorbance was read using ClarioStar Plus plate reader (BMG Labtech). The reciprocal endpoint titers were defined as the dilution with the OD_450nm_ 3 times higher than the background. Undetectable antibody titers (negative values) were assigned values of 0.

ELISA analysis of Spike protein exposure on EVs was performed as above, with few modifications. MaxiSorp ELISA plates were coated with S-EVs at 10 μg/ml in 100 μl 50 mM sodium carbonate/bicarbonate pH 9.6 buffer per well, overnight at 4 °C. Saturation was performed with 3% BSA in 1X PBS and followed by incubation with dilutions of either anti-S1 or S2 antibodies. Secondary HRP-conjugated donkey anti-rabbit and anti-mouse antibodies were used, respectively.

### IFN-γ ELISpot assay

Single cell suspensions were prepared from spleens of mice euthanized at day 63. Splenocytes from all immunized mice were analysed in two pools of three animals. Total T cells were isolated (EasySep™ Mouse T Cell Isolation Kit, StemCells #19851) and 2 × 10^5^ of T cells were stimulated for 18h in cell culture medium (RPMI 1640 with L-Glutamine, 25 mM Hepes, 10% iFBS and 5 μg/ml gentamycin) at 37 °C and 5% CO_2_ with overlapping peptide pools representative of either S1 or S2 SARS-CoV-2 proteins (JPT, #PM-WCPV-S) at 1 μg/ml each or DMSO in 96-well ELISpot IFN-γ plates (Mabtech #3321-4APW-2) in triplicates. As positive control stimuli, 0.5-1 × 10^5^ cells were stimulated with PHA-L (eBioscience, #15556286) at 1.25-5 μg/ml. Following the 18h incubation, plates were treated according to the manufacturer’s protocol. Acquisition and analysis were performed with a CTL Immunospot S6 analyser. Measurement values from DMSO stimulated cells were subtracted from all the measurements. Negative values were corrected to 0. Results are represented as a number of spot forming cells (SPC) per 1 × 10^6^ T cells.

### Pseudovirus neutralization assay

To produce SARS-CoV-2 pseudoviruses, SARS-CoV-2 truncated Spike expression and lentiviral pNL4-3.NanoLuc vectors were co-transfected into HEK293T cells using PEI. Pseudoviruses containing supernatant was collected 48h after, filtered through 0.45 μm filters, aliquoted and stored at −80 °C. To perform the neutralization assay 2-fold mouse sera dilutions (starting from 1:10) collected at day 63 were mixed with equal volumes of pre-titered (1000 times the background of fluorescence) Spike-pseudotyped HIV-NanoLuc viruses (1:2), incubated for 1h at 37 °C, 5% CO_2_ and then added to HEK293T cells transiently expressing ACE2 receptor and TMPRSS2 protease in 96-well plates (in triplicates) for 1h at 37 °C, 5% CO_2_. Cell culture medium was then changed. After a 48h incubation, cells were lysed and luciferase activity (RLU, relative light units) was measured using the Bright-Glo™ Luciferase Assay System (Promega). Background luminescence produced by cells-only controls (no pseudotyped virus) and positive luminescence produced by pseudotyped viruses infected cells were included and served to determine percentage of neutralization as 100% and 0%, respectively. Neutralization titers were defined as the sera dilutions that neutralize 50% of the virus.

### Protein sequence alignment

Multiple sequence alignment was performed with Clustal Omega tool (EMBL-EBI).

## Results

### EV-based vaccine design

Here, we present a SARS-CoV-2 vaccine candidate, CoVEVax which displays S1/S2 proteins naturally embedded in the membrane of EVs (hereafter, S-EVs) mimicking the original virus, with its MPER and TM sub-domains. Alignments of envelope sequences pointed out the importance and similarities of these two domains between coronaviruses and HIVs (Fig. 1, A and B). The HIV MPER required for the membrane fusion of HIV TM contains a tryptophan rich cluster, a cholesterol binding region and a neutralizing epitope conserved through all HIV clades (Fig. 1A) [14]. Interestingly, the S2 MPER sub-region of all coronaviruses (SARS-CoV-2, SARS-CoV, MERS-CoV and four human “common cold” coronaviruses) possesses an analogous highly conserved tryptophan cluster and a putative cholesterol binding region (Fig. 1B, top). Similarly to HIV, this conserved S2 MPER would contain a neutralizing B-epitope and it includes one of the T-cell epitopes cross-reactive in unexposed individuals [26]. In addition, the most reactive T-cell epitope of the SARS-CoV, entirely conserved in SARS-CoV-2, is located in the transmembrane domain (Fig. 1B, top) [27]. Altogether, these facts demonstrate the importance of including the MPER + TM domains in a vaccine.

**Fig. 1.**
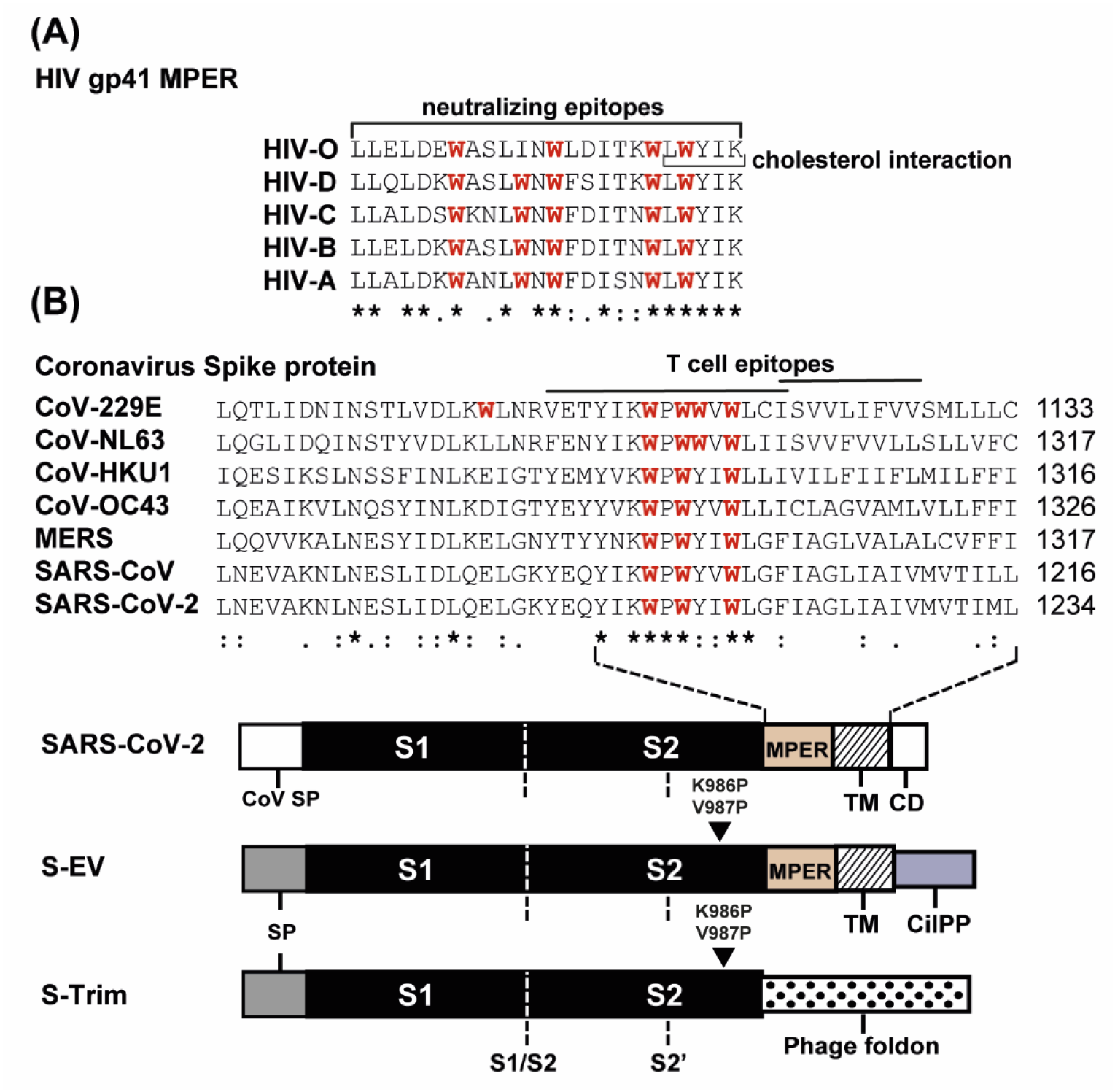
Alignments and predicted B- and T-cell epitopes within HIV gp41, coronavirus spike proteins, and tested vaccine candidates. (A) Alignment between HIV gp41 MPER sequences from different clades. A tryptophan (in red) rich region and cholesterol interaction domain are highly conserved among all HIV clades; MPER represents a conserved target of neutralizing antibodies. (B) Alignment of Spike MPER and TM domains of coronaviruses and constructs used in this study. A Tryptophan (in red) rich region and published T-cell epitopes localize in these domains. Both MPER and TM domains are retained in the Spike protein presented by S-EVs. These domains are deleted in the S-Trim and replaced by a phage trimerization motif merged downstream of Q1208 (bottom). S-EV and S-Trim contain at the N-terminus the Glut5 receptor SP; a pilot peptide (CilPP) addressing the Spike protein to EVs is fused at the end of S-EV TM domain. K986P, V987P: di-proline mutations. MPER: Membrane Proximal External Region; SP: Signal Peptide; TM: Transmembrane Domain; CD: Cytoplasmic Domain; CilPP: Ciloa Pilot Peptide; S-EV: Extracellular Vesicle carrying SARS-CoV-2 Spike at their surface; S-Trim: recombinant Spike protein Trimer.

Our first molecular construct and control immunogen, which was often used in other vaccines, comprises the full S1 protein, the S2 interrupted at the Glutamine 1208 and an additional phage foldon which allows the artificial trimerization (S-Trim, Fig. 1B and fig. 2A) [22]. This S1/S2 is soluble because it is devoid of the MPER and TM domains (Fig. 1B, bottom). The second construct, the basis of CoVEVax, is a S1/S2 anchored in a natural membrane, containing the MPER and TM domains identified as potentially important in triggering neutralizing humoral and cellular immune responses, fused to the proprietary Pilot Peptide (CilPP) which sorts the spike protein onto EVs (Fig. 1B, bottom).

**Fig. 2.**
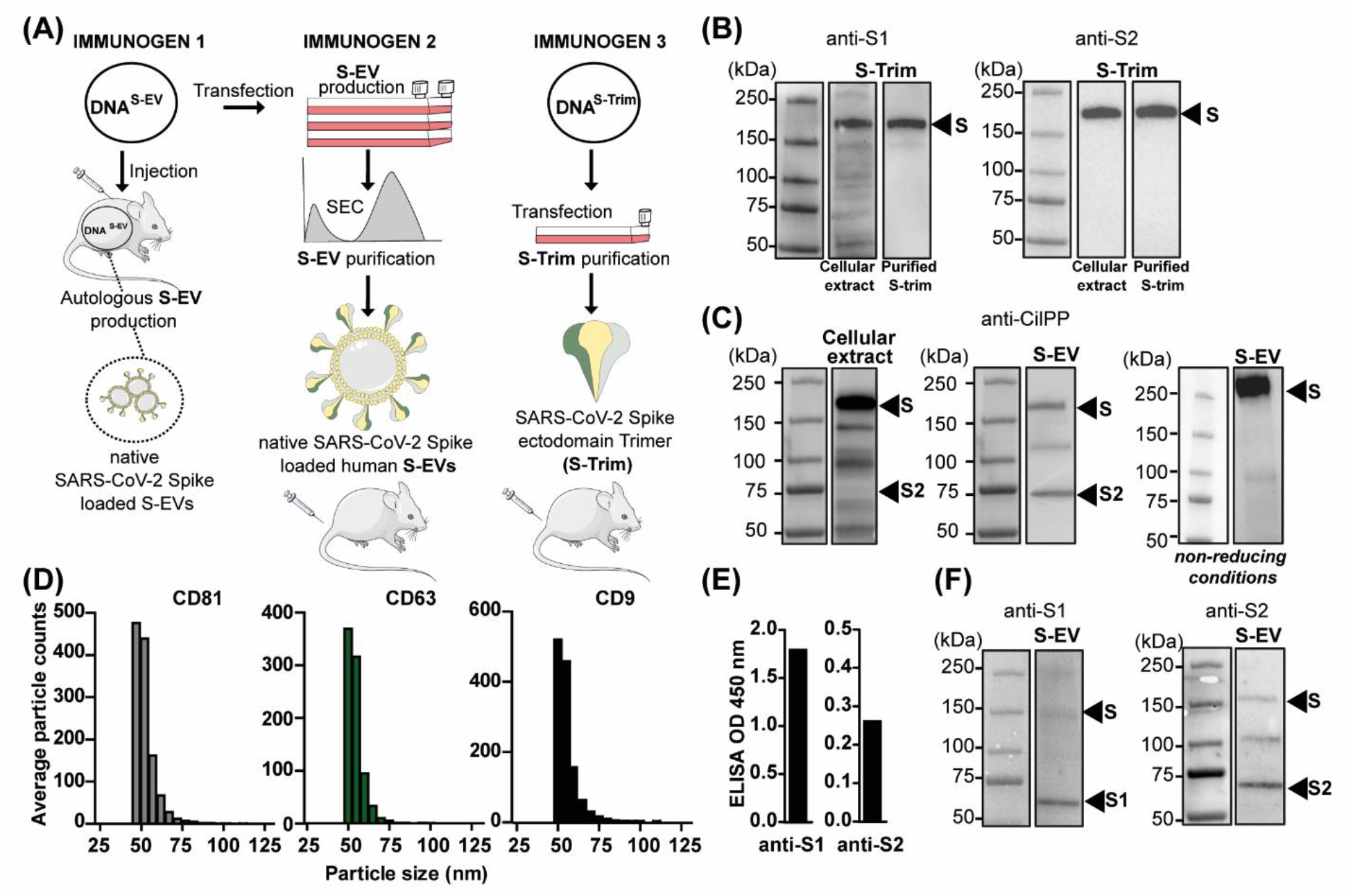
Production and characterization of the CoVEVax vaccine candidates. (A) Three different immunogens were generated to mount immune response against SARS-CoV-2 in mice. DNA^S-EV^ vector allows *in situ* production of autologous S loaded EVs upon injection into mice. S-EVs and S-Trim are produced in mammalian cells (HEK293T) *in vitro.* (B) WB analysis of purified SARS-CoV-2 S-Trim. (C) WB analysis of SARS-CoV-2 protein S expression in cellular extract and S-EVs in reducing (left and central panels) and non-reducing conditions (right panel) probed with anti-CilPP antibody. (D) Purified EV characterization by ExoView platform. EVs expressing SARS-CoV-2 S protein captured with either CD81, CD63 or CD9. (E) ELISA analysis of SARS-CoV-2 protein S1 and S2 presented on S-EVs; results present OD_450nm_ values. (F) WB analysis of SARS-CoV-2 protein S expression on S-EVs. DNA^S-EV^: DNA construct allowing production of SARS-CoV-2 Spike harbouring EVs; DNA^S-Trim^: DNA construct allowing production of S-Trim; SEC: Size Exclusion Chromatography.

### Production and characterization of CoVEVax

It is well-known that T-cell immune responses can be elicited by DNA vaccines, but that a protein boost is required for high humoral responses. Thus, we chose to develop a two component vaccine, the first one being a DNA^S-EV^ leading to *in situ* production and loading of S on autologous EVs and the second one, exogenously produced and purified S-EVs (Fig. 2A). As a control, a protein boost of spike protein trimer (S Trim) was also evaluated. The expression of S-Trim was analysed by Western blotting (WB). A band of the size expected for a non-mature protein was revealed when probed with either anti-S1 or anti-S2 antibody (Fig. 2B), indicating that the artificial S-Trim was correctly expressed but remained immature and was not processed into S1 and S2. DNA^S-EV^, when transfected in HEK293T cells allows the expression of two main proteins detected by an anti-CilPP antibody, the first with a size expected for an immature S and the second with a size expected for a mature S2 protein (Fig. 2C, left). This partial maturation of envelope proteins in cells was previously described for coronaviruses and HIV [28, 29]. Secreted EVs were purified and analysed for their size and protein content. Using the ExoView platform, we observed that S-EVs are specifically captured by anti-CD81, anti-CD63 and anti-CD9 antibodies and exhibit a size of ~55 nm, all being signatures of EVs (Fig. 2D). Enzyme-linked immunosorbent assay (ELISA) analysis revealed that these purified S-EVs harbour both S1 and S2 proteins at their surface (Fig. 2E). When analysed by WB, the secreted EVs contain immature S protein (Fig. 2C, middle and fig. 2F), predominantly mature S1 and S2 proteins when probed with anti-S1 or anti-S2 antibody, respectively (Fig. 2F); all being associated as a S1/S2 hetero-trimer naturally anchored in the EV membrane (Fig. 2C, right, non-reducing conditions).

### Study design and immune response against CoVEVax

Mice were immunized at day 0, 21 and 42 with 3 different combinations of described CoVEVax immunogens (Fig. 3A). Notably, in this study, no adjuvants were administered. Sera were analysed at day 42 (prime bleed) and 63 (terminal bleed) together with spleen collection (Fig. 3A). The humoral immune response induced by CoVEVax was assessed by ELISA. To determine SARS-CoV-2 specific antibody titers, we coated the ELISA plates with peptide pools representative of either the S1 or S2 protein. Compared to the negative control group 4 (PBS), all immunizations revealed evidence of antibodies directed against both S1 and S2 proteins after the protein boost. The mean titers against the S1 protein obtained with sera of group 1 (DNA^S-EV^/S-EV), group 2 (DNA^S-EV^/S-Trim) and group 3 (S-EV/S-EV) were 1716, 663, and 4316, respectively. The mean titers against the S2 protein obtained with sera of group 1, group 2 and group 3 were 1494,591,3116, respectively (Fig. 3B). The lowest titers were obtained with S-Trim boost injection and no S-EV injection, average titers were obtained with one S-EV boost injection and the highest resulted from three S-EV injections suggesting that titers correlate with the number of S-EV injections. Accordingly, sera from day 42 showed elevated ELISA titers within group 3 only (Fig. S1). Noteworthy, the humoral response against both S1 and S2 corresponding peptide pools is of a similar level, even though the S1 “head” with its RBD is widely recognized as the main immunogen. Here, when presented on EVs, the S2 “stem” acts as a potent antigen as well. In the depicted ELISA experiments only the humoral response against linear epitopes was measured. We hypothesized that the Spike presented on EVs would also trigger an immune response against conformational epitopes that are exhibited by the virus, which are generally the true targets of neutralizing antibodies. To this end, the neutralizing activity of the sera was quantified with a commonly used pseudotyped lentivirus neutralization assay (Fig. 3C). In brief, serial dilutions of sera were mixed with S-pseudotyped viruses and then used to infect HEK293T cells expressing human ACE2 receptor and TMPRSS2 protease. Results show that although the group 3 displayed highest ELISA titers, none of the animals generated neutralizing antibodies, similarly to the negative control group. In contrast, mice in group 2 have produced a good level of neutralizing antibodies (titers ranging from 320 to 2240, mean titer 973), which is in the range of the mean neutralizing titers reported for other vaccines [30-32]. Sustaining our hypothesis, sera of mice from group 1 also exhibited a very high level of neutralizing antibodies (titers ranging from 480 to 2430, mean titer 1472) (Fig. 3C). Finally, we looked at the T-cell mediated immunity, a second type of protective immune response, using ELISpot assay measuring antigen specific IFN-γ production. As before, we aimed to identify and discriminate between the immune responses against S1 and S2 proteins. Mice immunized with S-EVs only (group 3) did not develop cellular immune response as measured by IFN-γ ELISpot. By contrast, the two groups (1 and 2) immunized by two DNA^S-EV^ primes and one protein boost (either S-EV or S-Trim) developed strong cellular immune response, reaching ~350 spot forming cells (SFC) and ~150 SFC per 10^6^ T cells for S1 and S2 immunogens, respectively (Fig. 3D).

**Fig. 3.**
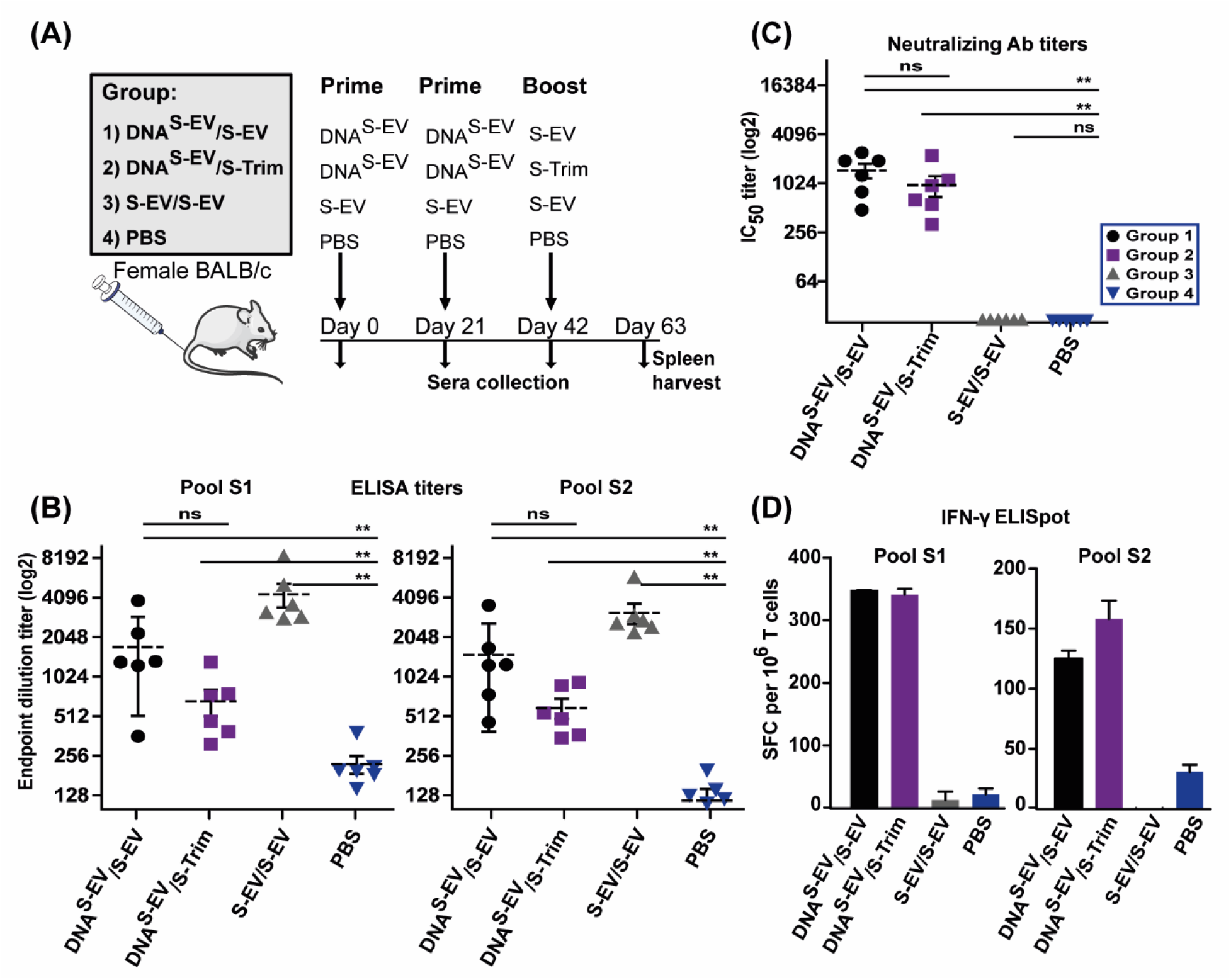
Humoral and cellular immune responses in mice immunized with CoVEVax. (A) CoVEVax immunization study design. Four groups of female BALB/c mice (n=6) received two prime injections at day 0 and 21 and a boost injection at day 42 according to the scheme. Mice were bled at day 0, 21, 42 (prime bleed) and 63 (boost bleed) and spleens were harvested. (B) Specific IgG response to SARS-CoV-2 S1 or S2 peptide pools at day 63 measured by ELISA. Results are presented as reciprocal endpoint dilution titers. Data present mean ± SEM. Dashed lines indicate mean titers. (C) Pseudovirus neutralizing antibody titers of sera collected at day 63. Titers correspond to the sera dilutions neutralizing 50% of the virus. Data present mean ± SEM of two assay replicates (with each serum sample tested in triplicates). Dashed lines indicate mean titers. (D) IFN-γ ELISpot analysis of antigen specific cellular response. Total T cells were isolated from pooled splenocytes and stimulated with either S1 or S2 SARS-CoV-2 peptide pools. Results are presented as the number of spot forming cells (SPC) per 1 × 10^6^ T cells. **p<0.01, calculated by Mann-Whitney test.

## Discussion

In conclusion, we have developed a unique EV-based vaccine platform likely adaptable to several enveloped viruses. Here, we establish its proof of concept with EVs harbouring a fully native and mature Spike of the SARS-CoV-2. The fact that most of the S proteins are matured into S1 and S2 and that they remain associated as hetero-trimers pleads for a fully native conformation of the Spike presented by EVs. It appears that keeping the S transmembrane domain and merging it with the CilPP allows to obtain this fully native conformation. This contrasts with a previous SARS-CoV EV-based construct where the S protein remained immature, perhaps because the transmembrane and the cytoplasmic domains of S were exchanged for those of the VSV-G protein [6]. During our vaccine design we have preserved specific envelope protein domains (MPER and TM) often omitted in other SARS-CoV-2 vaccine candidates, which proved crucial in mounting immune responses against other viruses. CoVEVax triggered strong neutralizing and cellular immune responses. These responses were obtained in the absence of any other viral components and without adjuvants or other macromolecules. Many vaccine approaches targeting SARS-CoV-2 are based on viral vectors, particularly adenoviruses, including the most immunogenic ones from human serotype group C (Ad5) [33]. Despite the fact that they are well characterized, it is clear now that one of their major limitations is the natural, pre-existing immunity against these vectors [33, 34]. Likely, all primarily vaccinated hosts will mount an anti-adenovirus immune response that will hamper a strong anti-SARS-CoV-2 response after a second injection. As a consequence, adenoviral vectors from other species or with a lower seroprevalence in humans must be explored as vaccine carriers [30, 32, 35].

One of the major concerns with emerging viruses is the threat of re-infections in the general population lacking a long-lasting immunity contributing to the virus becoming endemic, which in turn may be linked with increased mutational-rate, as in the case of influenza virus, necessitating yearly vaccination campaigns. Due to the high immunogenicity of viral vectors themselves multiple immunizations with such platforms could prove challenging, if not inefficient. EV antigenic display can overcome this problem. Moreover, it has been demonstrated that an antigen embedded in the membrane of EVs is more immunogenic and triggers a higher humoral response *in vivo* when compared with human (Ad5) and non-human (ChAdOx1) adenoviral vectors, both proposed as vaccine platforms against SARS-CoV-2 [36]. The immune responses observed upon CoVEVax administration *in vivo* are directed against both S1 an S2 parts of the Spike protein, which is important for a broad coverage of the vaccine in regard to the high conservation of the S2 protein through all coronaviruses clades. The combination of two components in the CoVEVax is crucial to obtain both types of protective immune responses, the DNA^S-EV^ based immunization triggering T-cell response, while the S-EV immunization rising the humoral response. The DNA components of this vaccine platform are of high importance also because the released EVs express S1 and S2 proteins in an environment of only fully autologous proteins, contributing to the safety of the prime injections. The relatively low global binding antibody levels associated with a strong neutralizing activity prompt us to expect that CoVEVax is unlikely to promote the induction of antibodies which can trigger Antibody Dependent Enhancement (ADE) adverse events. This EV-based vaccine which contains highly conserved S2 sequences, will in all likelihood exhibit broader protection even against SARS-CoV-2 variants that will began to appear by antigenic drift [37, 38]. It is likely that in contrast to soluble protein vaccines, but similarly to viral ones, EVs containing multiple copies of membrane-anchored S proteins should better mimic the SARS-CoV-2 virions. Exposure of several S-proteins in their natural conformation on the surface of the same EV should facilitate B-cell receptor crosslinking. In addition, the DNA injection allows S presentation in association with autologous cellular components that might provide additional immunostimulatory signals. Importantly, EV-based vaccines have first raised interest because they permit efficient CD8+ T-cell responses. Our results sustain entirely all these hypotheses.

Finally, the characteristics of our EV-based vaccine platform should prove an important countermeasure in the ongoing COVID-19 pandemic but also facilitate quick development of future vaccine candidates against emerging viral threats.

## Acknowledgments

The authors thank Christine Chable-Bessia at CEMIPAI CNRS UMS 3725 − University of Montpellier, Montpellier, France, for technical assistance in BSL3 facilities. We thank the animal staff from Metamus facility which belongs to Montpellier animal facilities network (RAM, Biocampus). We thank Dr. Clement Mettling for providing the pNL4-3.Luc.R-E-luciferase reporter vector.

## Additional information

## Author contributions

KP, BT and RZM designed research; KP, NG, ML, DM and SC performed research; CB-G contributed to immunizations; KP, BT and RZM wrote the manuscript.

## List of Supplementary Materials

**Fig. S1**

