## Supplementary Materials for "Extracellular vesicle-based vaccine platform displaying native viral envelope proteins elicits a robust anti-SARS-CoV-2 response in mice"

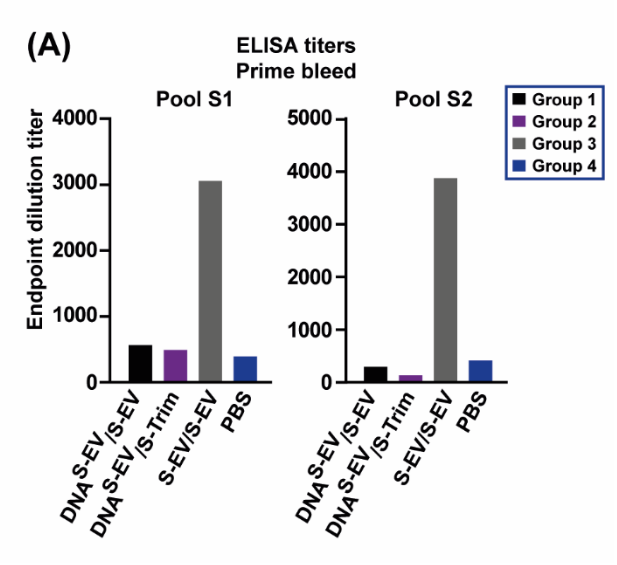


**Fig. S1. Humoral response in mice immunized with CoVEVax.**

(A) Specific IgG response to SARS-CoV-2 S1 and S2 peptide pools at day 42 measured by ELISA. Results are presented as reciprocal endpoint dilution titers from pooled sera; n=6 mice per group.
